# SARS-CoV-2 infection in domestic rats after transmission from their infected owner

**DOI:** 10.1101/2022.10.13.512053

**Authors:** Xavier Montagutelli, Bérénice Decaudin, Maxime Beretta, Hugo Mouquet, Etienne Simon-Lorière

**Author notes:** corresponding author, Mouse Genetics Laboratory, Institut Pasteur, 25 rue du Dr Roux, 75724 Paris cedex 15, France.

## Abstract

We report the transmission of SARS-CoV-2 Omicron variant from a COVID-19 symptomatic individual to two domestic rats, one of which developed severe symptoms. Omicron carries several mutations which permit rodent infection. This report demonstrates that pet, and likely wild, rodents could therefore contribute to SARS-CoV-2 spread and evolution.

## Main text

In May 2022 (week 18), a resident of the Southern outskirts of Paris, France, without recent history of travel, developed flu-like symptoms with moderate fever, headache, coughing, shortness of breath and fatigue, which resolved after 8 days. A SARS-CoV-2 antigenic test was positive three days after the onset of symptoms (day 3) and was confirmed by a RT-qPCR test performed on day 5. On day 12, one of the owner’s two pet rats (Rat 1), a 3-year-old male, developed prostration, respiratory distress (open-mouth breathing, cyanosis, chromodacryorrhea, pulmonary cracks) and anorexia. This rat had a history of frequent sneezing which had worsened during the previous weeks. The two rats had been in close contact with their owner for cuddles. On day 12, Rat 1 was presented to a veterinarian who noted very severe symptoms (respiratory distress and lethargy) responsible for marked suffering. These symptoms worsened three days after admission, leading to the decision to euthanize the rat on D15. A blood sample was collected, but no other samples were available for further analysis. On day 26, the owner presented to the veterinarian Rat 2, a 6-month-old male, which was in good condition with only occasional sneezing. A blood sample was collected for serology.

Infection of the owner with a SARS-CoV-2 variant of the Omicron lineage was evidenced by the detection of the K417N mutation in the PCR test performed 5 days after symptoms offset (week 18 of 2022), and compatible with the 100% proportion of Omicron reported by Santé Publique France in the sequencing of randomly selected SARS-CoV-2 positive samples in France during week 17 and 18 of 2022. Data from the National Reference Center for the Northern part of France further indicate that Omicron BA.2 was dominant at this time in the Paris area.

The sera of both pet rats presented IgG and IgM titers against both Wuhan and Omicron spike antigens when assayed by ELISA as previously described (Montagutelli, 2021, Planchais, 2022), in parallel to control sera collected from two healthy rats (Rat 3 and Rat 4) bred under specific-pathogen-free condition (Figure 1). This result indicates that the pet rats had been recently exposed to SARS-CoV-2. While we cannot entirely exclude a more distant infection, considering that the two rats were kept in a cage inside the owner’s home, away from other animals, their contamination can most likely be attributed to their close and frequent contacts with the infected owner in May 2022.

**Figure 1.**
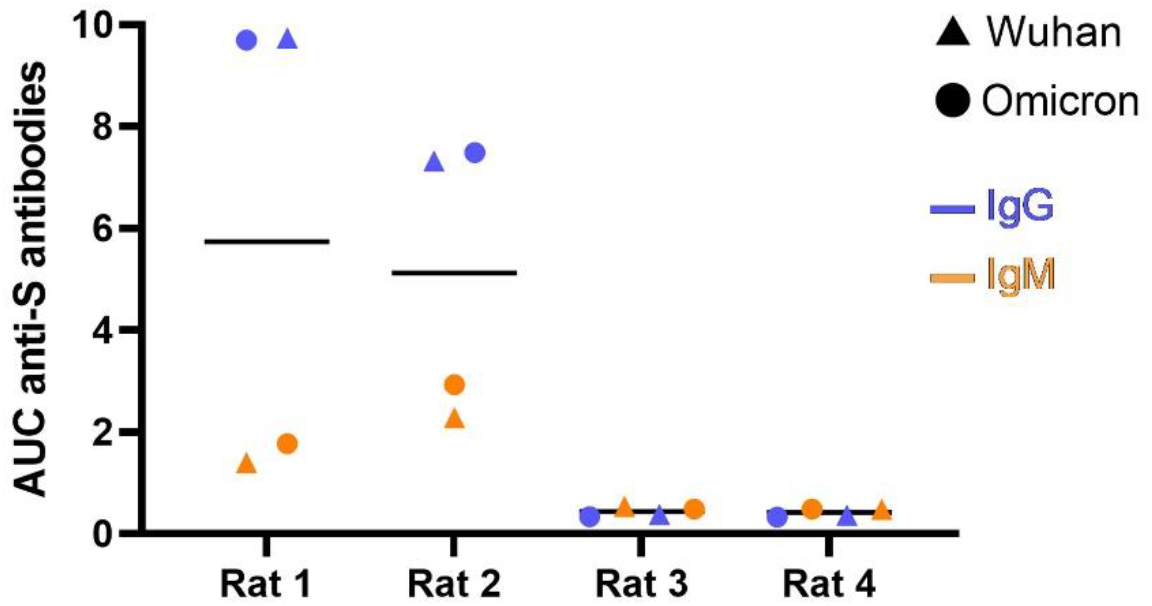
Serum anti-SARS-CoV-2 spike IgM and IgG antibody titers in rats. ELISA antibody reactivities against Wuhan and Omicron spike antigens as shown as area under the curve (AUC) of optical density values from serum titrations. Rat 1 developed a severe condition and was euthanized 3 days after symptoms onset. Rat 2 was minimally affected. Rat 3 and Rat 4 were two uninfected rats. Rats 1 and 2 presented very high IgG titers recognizing both spikes.

While the initial SARS-CoV-2 variants were shown to be unable to infect rodents, this changed with the evolution of the virus during the pandemic circulation in humans. In particular, multiple Variants of Concern have been shown to be able to infect rodents in experimental settings. This extension of the host range has been associated with changes in the spike protein such as the recurrent N501Y mutation (Montagutelli, 2021). Omicron variants have been shown to readily infect laboratory rodents, with varying disease severity, and this capacity and variations likely involve other changes in parallel to N501Y (Montagutelli, 2022).

Human-to-animal SARS-CoV-2 transmission has been reported in several domestic animal species including ferrets (Gortázar, 2021), minks (Oude Munnink, 2021), rabbits (Fritz, 2022), cats (Klaus, 2021) and hamsters (Yen, 2022), as well as in free-ranging white-tailed deer populations in North America (Hale, 2022). Although mice and rats are occasionally kept as pet animals, they mostly live as wild animals, and many species are human commensals. This report shows that human to rat infection can occur in a natural setting. In addition, the timing of clinical signs is compatible with transmission between Rat 1 and Rat 2, which was shown to happen in mice by close contact in the same cage (Montagutelli, 2021). The possibility that rodents may host SARS-CoV-2 and transmit the virus to congeners supports the hypothesis that wild rodents may become secondary reservoirs of SARS-CoV-2, where the virus may evolve differently than in humans, and should be under epidemiological surveillance (Montagutelli, 2022). This hypothesis has been notably raised to explain the detection of cryptic SARS-CoV-2 lineages in wastewaters (Smyth, 2022). Nevertheless, pet rodent owners should refrain from being in contact with their animals during the PCR-positive phase of SARS-CoV-2 infection.

## Acknowledgements

We are grateful to the owner of the two rats for participating in this study. We thank Dr Charly Pignon at Ecole nationale vétérinaire d’Alfort for reaching out to veterinarians taking care of pet rats. We thank Nathalie Jolly and Frédéric Gouin of the Institut Pasteur Clinical Core for the collection of medical information

This project was supported by a grant from the French Government’s Investissement d’Avenir programme, Laboratoire d’Excellence “Integrative Biology of Emerging Infectious Diseases” (grant n°ANR-10-LABX-62-IBEID). ESL acknowledges funding from the INCEPTION programme (Investissements d’Avenir grant ANR-16-CONV-0005), from the PICREID program (Award Number U01AI151758)

## Ethics statement

The authors confirm that the ethical policies of the journal, as noted on the journal’s author guidelines page, have been adhered to. No ethical approval was required as all research has been done on diagnostic specimen.

## Author contribution

XM designed the study. BD collected clinical data and samples. MB and HM analyzed the samples. XM and ESL wrote the manuscript with input from all authors.

## Methods

### Anti-S ELISA

High-binding 96-well ELISA plates (Costar, Corning) were coated overnight with purified Wuhan and Omicron BA.1 SARS-CoV-2 trimeric Spike proteins (125 ng/well in PBS). After washes with 0.1% Tween 20 in PBS (PBST), plates were blocked for 2h with PBS added with 1% Tween 20, 5% sucrose and 3% milk powder (Blocking solution). After PBST washes, 1:50-diluted rat sera (in PBST + 1% BSA) and 7 consecutive 1:3 dilutions were added and incubated for 2 h. After washes, plates were revealed by addition of goat HRP-conjugated anti-rat IgM or IgG (0.8 μg/ml final in blocking solution, Immunology Jackson ImmunoReseach). Plates were revealed by adding 100 μl of HRP chromogenic substrate (ABTS solution, Euromedex) after PBST washes. Experiments were performed in duplicate at room temperature using a HydroSpeed microplate washer and Sunrise microplate absorbance reader (Tecan, Mannedorf), with OD at 405nm. Area under the curve (AUC) values were determined by plotting the log10 of the dilution factor values (X axis) required to obtain OD 405nm values (Y axis). AUC calculation analyses were performed using GraphPad Prism software (v8.4.1, GraphPad Prism Inc.).

